# Modeling Meibomian Gland Development and Dysfunction: A Mouse-Derived Organoid System Reveals Hippo-YAP as a Critical Regulator

**DOI:** 10.64898/2026.05.13.724874

**Authors:** Meiqin Zhong, Jingbin Zhuang, Lingyu Zhang, Rongrong Zhang, Le Sun, Wei Li, Yang Wu, Jinghua Bu

## Abstract

The developmental program governing meibomian gland (MG) morphogenesis and proliferation remains poorly understood, largely due to the lack of physiologically relevant model systems. Here, we established a novel high-fidelity, three-dimensional organoids model derived from mouse meibomian gland (mMGO) epithelium. Transcriptomic and phenotypic analyses demonstrated that mMGOs faithfully recapitulate postnatal gland development *in vivo*, including dynamic transcription program, branching morphogenesis, lineage differentiation, and functional lipid accumulation. Leveraging this model, we identified the Hippo-YAP pathway as a pivotal regulator of MG epithelial proliferation and homeostasis for the first time. YAP inhibition severely impaired organoids growth, while pharmacological inhibition of Hippo pathway with XMU-MP-1 enhanced proliferation and progenitor clonogenicity. Crucially, in inflammation-induced atrophic organoids, XMU-MP-1 treatment rescued YAP nuclear localization and stimulated regrowth and functional restoration. Our study provided new mechanistic insights and a robust organoids platform for MG development research, and nominated targeted Hippo pathway inhibition as a promising strategy to reverse glandular atrophy in meibomian gland dysfunction.

## 1. Introduction

Meibomian gland (MG), a specialized sebaceous gland embedded in the eyelid, prevents tear evaporation and maintains ocular surface homeostasis by secreting the lipid layer of the tear film.^[1]^ Meibomian gland dysfunction (MGD) is the leading cause of evaporative dry eye disease which affects hundreds of millions globally and severely compromising quality of life.^[2, 3]^ Clinically, MGD is characterized by ductal obstruction, altered lipid secretion, and progressive glandular atrophy, ultimately resulting in chronic ocular surface inflammation and visual disturbance.^[1]^ Despite its high prevalence and clinical burden, the underlying mechanisms governing MG development, homeostasis, and regeneration remain incompletely understood. A major obstacle in elucidating MG biology lies in the lack of physiologically relevant experiment models.^[4]^

Traditional *in vivo* animal models provide valuable insights but are limited by complexity, low throughput, and difficulty in dissecting cell-type-specific mechanisms. Meanwhile, conventional two-dimensional (2D) primary culture fails to recapitulate the three-dimensional (3D) architecture and cell-cell interactions essential for gland morphogenesis and function.^[4, 5]^ Recent advances in organoid technology have enabled the establishment of self-organizing, stem cell derived 3D cultures that closely mimic native tissue structure and function. Organoid systems have been successfully applied to multiple epithelial organs, including intestine,^[6]^ liver,^[7]^ and skin appendages.^[8]^ Recent progress has led to the generation of 3D organoid systems from both mouse and human MGs.^[9, 10]^ These organoids replicated some key tissue features and have been used for disease modeling and drug testing of known pathways. However, whether MG organoid models can be effectively leveraged to investigate MG development, and more importantly, whether they can uncover the key regulatory mechanisms governing this process, remains to be determined.

In this study, we developed a novel murine MG organoid system, which closely mimicked the dynamic morphogenetic and transcriptional timeline of postnatal development. Utilizing this platform, we identified the Hippo-YAP pathway as a previously unreported crucial regulator of MG epithelial proliferation and homeostasis.

Furthermore, we demonstrate that pharmacological inhibition of this pathway can reverse established organoids atrophy and reactivate growth. Our study thus not only complements existing models by providing both a platform for studying MG organogenesis, but also uncovers a key regulatory mechanism governing glandular development. These findings offer new insights into the pathophysiology of MGD and may inform the development of regenerative and targeted therapeutic strategies.

## 2. Results

### 2.1 Establishment and characterization of mouse meibomian gland organoids

To establish an *in vivo* benchmark, we characterized postnatal mouse MG development from day 4 postnatal (P4) to P60. *In vivo* confocal imaging revealed progressive branching morphogenesis, forming a complex acinar network by P60 (Figure S1A, Supporting Information). Immunofluorescence showed sustained expression of the basal/progenitor marker Keratin 14 (K14), and the ductal differentiation marker Keratin 6a (K6a) depicted the developing ductal system (Figure S1C, Supporting Information).^[11]^ The lipidogenic regulator PPARγ exhibited nuclear translocation by P10, marking the acinar cell maturation (Figure S1B, Supporting Information), while LipidTox and Oil Red O staining demonstrated progressive neutral lipid accumulation from P4 to P14 (Figure S1B, D, Supporting Information). These data define a timed developmental program as an *in vivo* benchmark.

Then, we sought to generate mouse meibomian gland organoids (mMGOs) that recapitulate this morphogenesis and molecular features *in vitro*. Meibomian gland cells (MGC) and 3T3-J2 cells were co-cultured in hemispherical matrigel with the screened optimal concentration of matrigel and cell proportion (Figure S2A,B, Supporting Information).

The mMGOs recapitulated key developmental stages *in vitro*. Morphologically, they progressed from solid spheroids (D4) to form branched, multi-acinar structures with clear lumens by day 10, as shown by brightfield, H&E staining (Figure 1, A,B). Molecular characterization confirmed the expression of canonical MG lineage markers, including K14 (basal/progenitor cells), K6a (ductal epithelial cells), and K10 (keratinized epithelial cells) (Figure 1C). Proliferation, marked by Ki67, was localized to budding regions (Figure 1C). Critically, within the innermost region of the organoids, we observed LipidTox-positive acinar cells. Surrounding these, multiple layers of K6a-positive ductal epithelial cells were present. The outermost layer comprised a small number of K14-positive, weakly K6a-positive transitional cells, further confirming the cellular differentiation within the mMGOs (Figure 1D). Additionally, the organoids exhibited functional maturation, with significant nuclear staining of the lipidogenic regulator PPARγ by day 7 and concomitant accumulation of LipidTox-positive neutral lipids from day10 onward (Figure 1E). Overall, the organoids recapitulate the morphology, cellular lineage, and lipid synthesis capacity of the native MG, establishing a faithful *in vitro* system.

**Figure 1.**
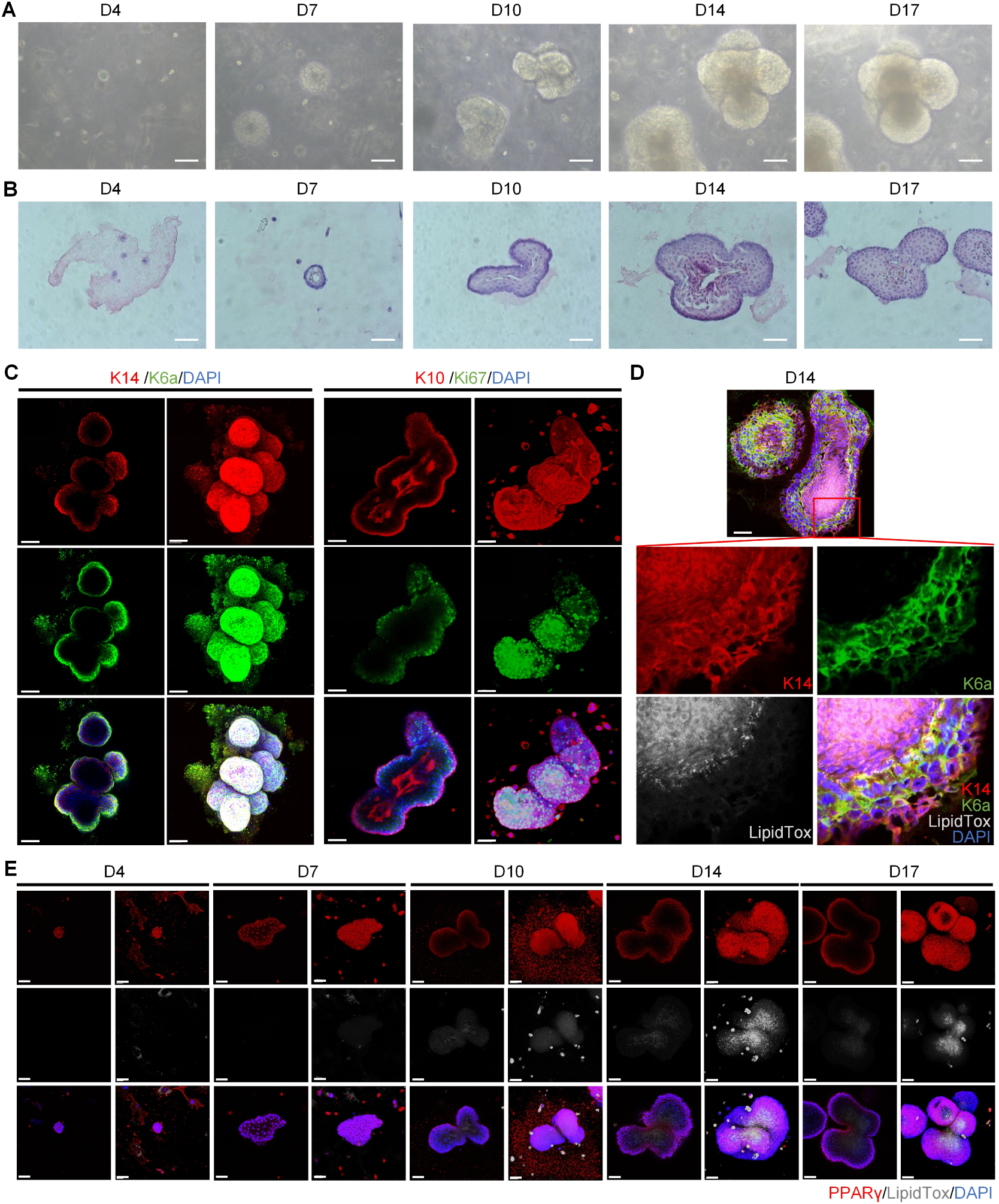
Characterization of mMGOs. (A) Bright-field images of mMGOs during formation. (B) Representative H&E images of mMGOs during formation. (C) Representative immunofluorescent images of K14, K6a, K10 and Ki67 of mMGOs on day 10. (D) LipidTox staining and immunofluorescence of K14 and K6a of mMGOs section on day 10. (E) LipidTox staining and immunofluorescence of PPARγ of mMGOs during formation. Scale bars represent 80 µm in (A), 50 µm (C), (D) and (E), 40 µm in (B).

### 2.2 Transcriptomic profiling reveals mMGOs recapitulate developmental and functional programs of the mouse MG

To molecularly evaluate the fidelity of our mMGOs, we performed RNA sequencing across multiple developmental time points of both mouse MG *in vivo* and organoids *in vitro*. Sample correlation analysis revealed a strong temporal concordance, with early organoid stages (D4, D7) clustering closely with early postnatal glands (P4, P7), and later organoid time points (D9, D12 and D15) aligning with more mature gland stages (P10, P12 and P14) (Figure 2A). This developmental congruence was further substantiated by Procrustes analysis, demonstrating a significant overall similarity between the transcriptional landscapes of the mMGOs and mMG datasets (M² = 0.5226, p = 0.001) (Figure 2B). Gene Ontology (GO) enrichment analysis indicated that mMGOs recapitulate the functional programs of *in vivo* MG development in a stage-specific manner. During the early phase (mMGO D4-D7 versus *in vivo* P4-P7), both systems showed robust enrichment for core developmental processes (Figure 2C), including system development, anatomical structure morphogenesis, and tissue development, etc. During the subsequent phase (mMGO D7-D9 versus *in vivo* P7-P10 and P10-P12), the two systems continued to exhibit concordant enrichment, such as lipid metabolic process, response to external biotic stimulus, and response to other organism (Figure 2C). Collectively, these comparative transcriptomic data establish that mMGOs undergo a developmental program that closely mirrors the temporal and functional progression of the native MG.

**Figure 2.**
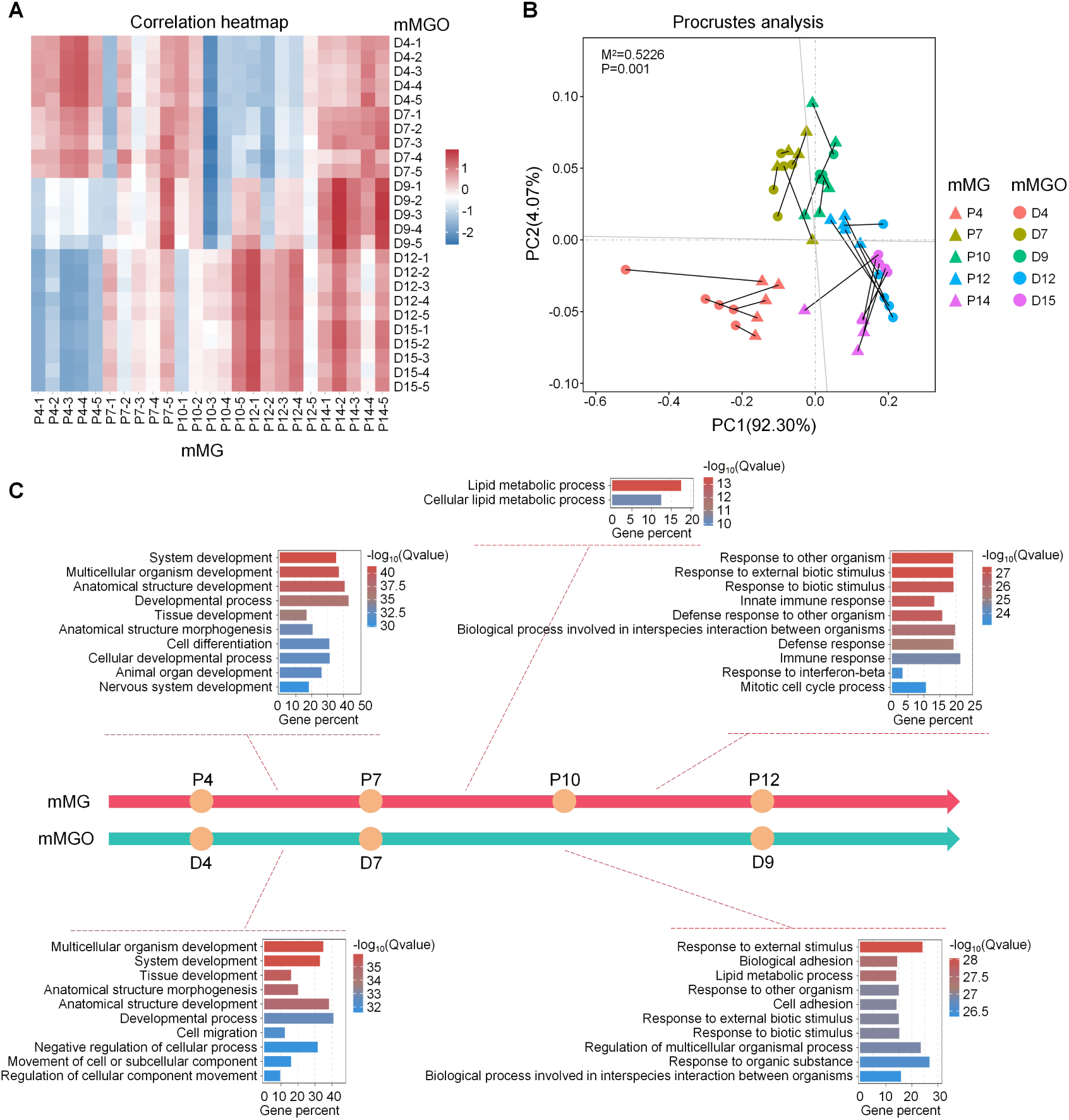
Comparative transcriptomics demonstrates high similarity between mMGOs and native gland development. (A) Correlation heatmap of mMGOs and mouse MG at multiple time points. (B) Procrustes analysis of mMGOs and mouse MG at multiple paired time points. (C) GO enrichment analysis of biological process for DEGs identified in mMGOs (D7 versus D4, D9 versus D7) and mouse MG (P7 versus P4, P10 versus P7, P12 versus P10). mMG: mouse meibomian gland.

### 2.3 Activation of Hippo-YAP pathway during mMGOs and mouse MG development

By the day 7 of culture, the organoid volume had increased to several times that of day 4, and the budding had begun to appear (Figure 1A), indicating that day 7 represents a critical developmental milestone for organoids, marking the peak of both proliferation and morphogenesis. To investigate the key biological processes and their regulatory mechanisms during this particular stage, a Gene Set Enrichment Analysis (GSEA) was performed.

Compared to day 4, day 7 shows significant enrichment of spliceosome, DNA replication, nucleocytoplasmic transport, tight junction and cell cycle (Figure 3A). We also identified enrichment of the RIG-I-like receptor signaling pathway, Fanconi anemia pathway and Hippo signaling pathway (Figure 3A). Reactome enrichment analysis revealed that the genes regulations were predominantly associated with CDC42, CDK1 and AURKB (Figure 3B). Protein-Protein Interaction Networks (PPI) showed that the core proteins of the top 100 pairs of interactions include CDC20, TRAF6 and MCM7 (Figure 3C). Notably, these genes above contribute to the regulation of the cell cycle by the Hippo-YAP pathway.^[12]^ Subsequently, GSEA revealed a significant upregulation of Hippo signaling pathway (Figure 3D), and the heatmaps indicated that the downstream genes of Hippo-YAP pathway were upregulated on day 7 (Figure 3E). Moreover, the nuclear translocation of YAP was demonstrated during mMGOs (Figure 3F,G) and mouse MG development (Figure 3H,I). These data suggest that Hippo-YAP is the predominant signaling pathway significantly activated during the development of the mMGO and mouse MG.

**Figure 3.**
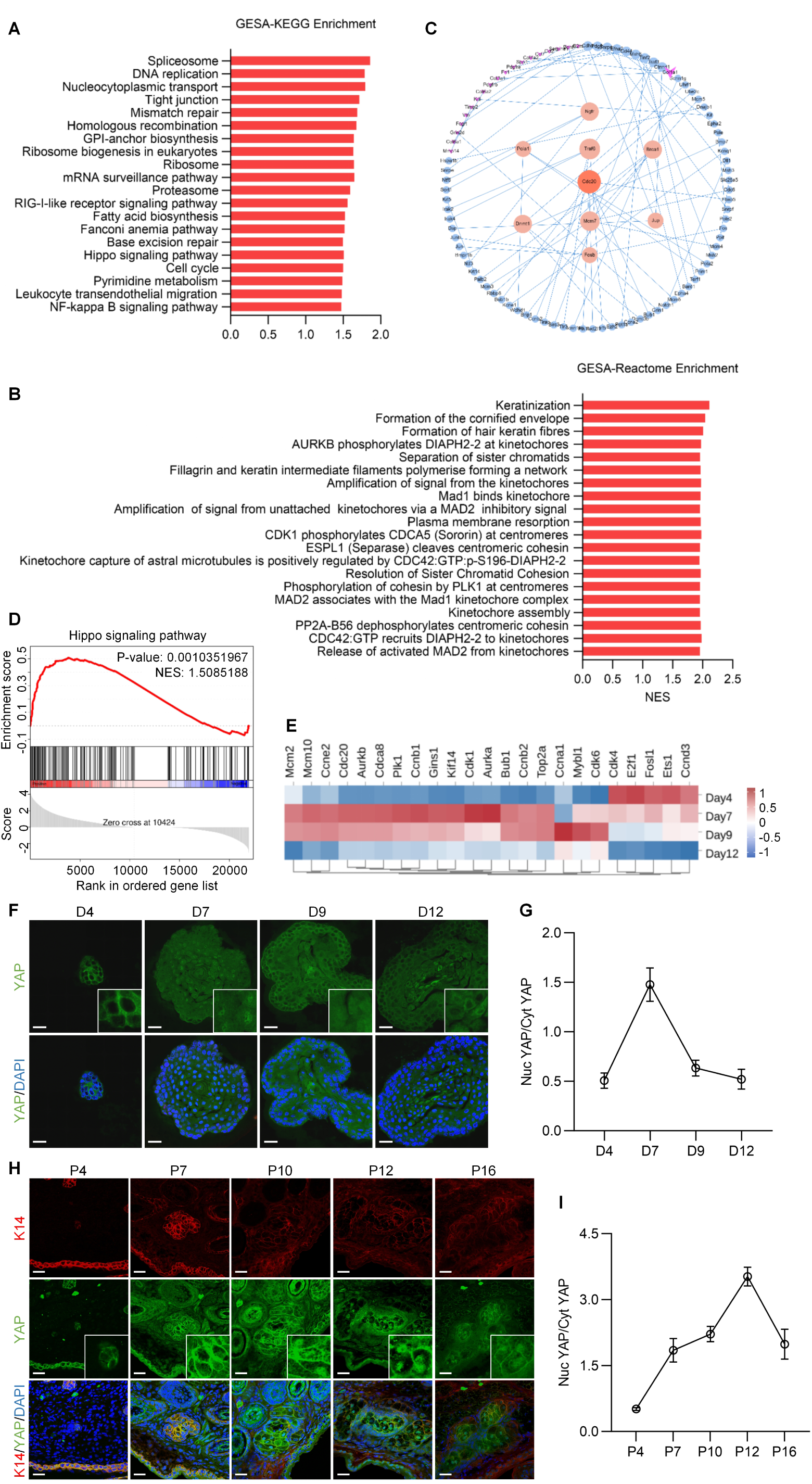
Hippo-YAP signaling pathway is activated during mMGO and mouse MG development. (A) Top 20 KEGG items with positive NES values from mMGOs GSEA for day7 versus day4. (B) Top 20 Reactome items with positive NES values from mMGOs GSEA for day7 versus day4. (C) Day7 versus day4 mMGOs PPI network analysis for the top 100 pairs of protein interactions. (D) The Hippo signaling pathway GSEA plot for mMGOs on day 7 versus day 4. (E) Heatmap of cell cycle-related genes downstream of the Hippo signaling pathway in mMGOs from day 4 to day 12. (F and G) Representative immunofluorescence images of YAP in mMGOs and the ratio of nuclear YAP to cytoplasm YAP, four cells were randomly selected for measurement and averaged in each sample (n=4). (H and I) Representative immunofluorescence images of YAP in mouse MG, positioned by K14. And the ratio of nuclear YAP to cytoplasm YAP (n = 3). Scale bars represent 25 µm in (F) and (H). Nuc, nuclear; Cyt, cytoplasmic.

### 2.4 YAP inhibition and knockdown of *Yap1* suppress mMGOs formation

We further investigated the function of Hippo-YAP pathway on mMGOs formation. Verteporfin, a specific YAP inhibitor disrupting the YAP-TEAD interaction,^[13]^ was used to treat mMGOs during early stage. As shown in Figure 4A,B, YAP inhibition markedly suppressed mMGOs formation, quantified by axis length and surface area. Immunofluorescence of Ki67 and EdU staining showed the cell proliferation of mMGOs was significantly inhibited (Figure 4C,D,E,F), which also supported by the relative mRNA expression levels of *Ki67* and *Pcna* (Figure 4G).

**Figure 4.**
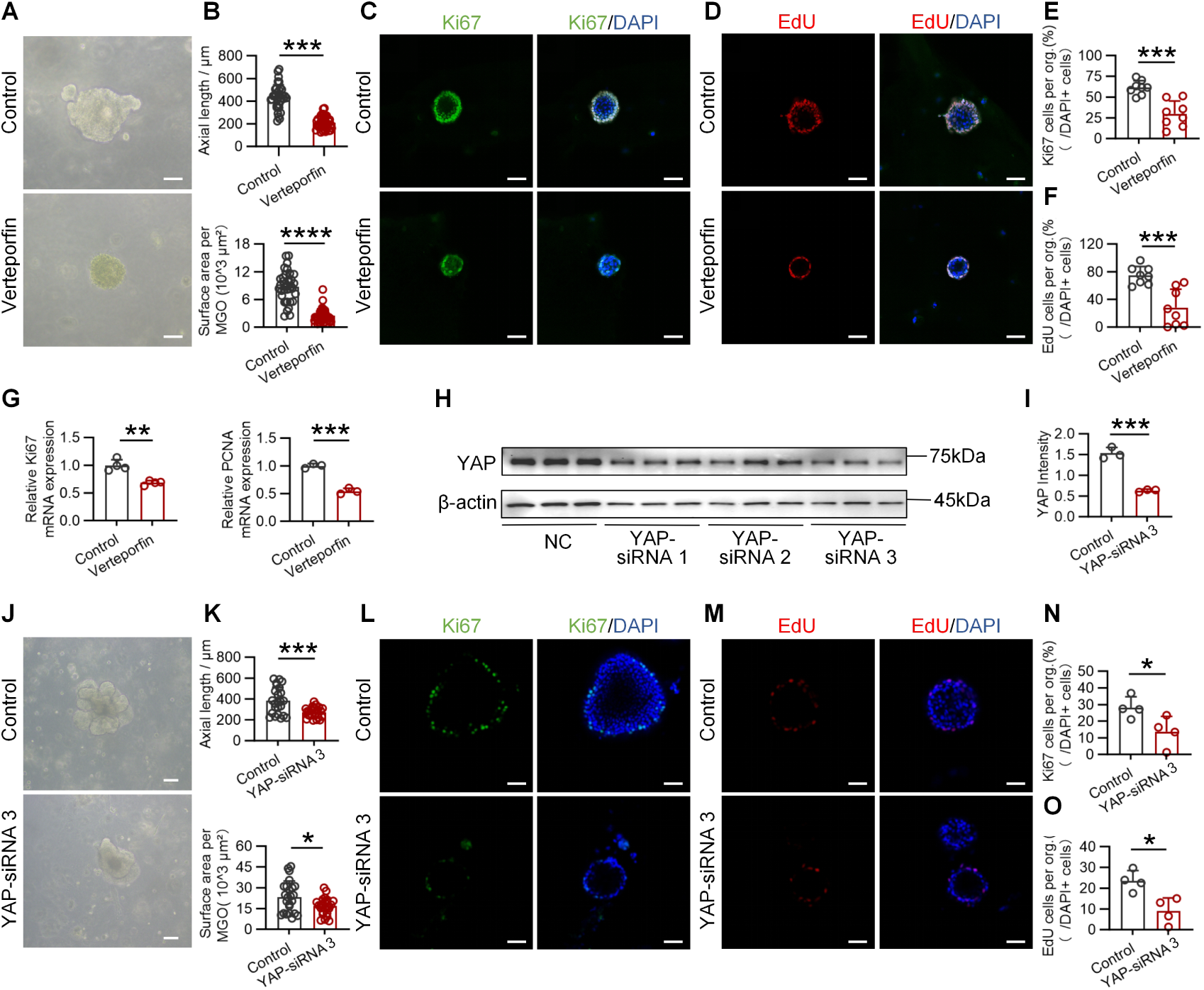
YAP inhibition and *Yap1* knockdown suppress mMGO formation and proliferation. (A and B) Bright-field images of mMGOs treated with verteporfin, with DMSO as control. The axial length and the surface area of mMGOs were measured using ImageJ (n=35). (C and D) Immunofluorescence of Ki67 and EdU staining in mMGOs, nuclei were counterstained with DAPI. (E and F) Percentage of Ki67 and EdU positive cells in (C and D) (n=8). (G) Relative mRNA expression of *Ki67* (n=4) and *Pcna* (n=3) in mMGOs. (H and I) Western blot analysis of YAP in mMGOs transfected with 3 types of YAP-siRNA, with NC-siRNA as control, the protein levels were quantified by densitometry (n=3). (J and K) Bright-field images of mMGOs transfected with YAP-siRNA 3, with NC-siRNA as control. And the axial length and the surface area of mMGOs were quantified (n=24). (L and M) Immunofluorescence of Ki67 and EdU staining in mMGOs, nuclei were counterstained with DAPI. (N and O) Percentage of Ki67 and EdU positive cells in (L and M) (n=4). Scale bars represent 100 µm in (A) and (J), 50 µm in (C), (D), (L) and (M).

In addition to pharmacological inhibition, a siRNA was used to konckdown *Yap1* in mMGOs. As shown by western blots, YAP expression was reduced after transfection (Figure 4H,I). YAP-siRNA group exhibited smaller sizes and fewer buds compared to the control (Figure 4J,K). EdU and Ki67 positive cells also significantly decreased after knockdown of *Yap1* (Figure 4L,M,N,O). Collectively, these results indicated that Hippo-YAP pathway plays a pivotal regulatory role during the initial stage of mMGOs formation.

### 2.5 Hippo pathway inhibition promotes mMGOs formation and enhances stem/progenitor cells stemness

YAP is directly activated by the Hippo pathway inhibition.^[14, 15]^ To investigate the effects of YAP activation on mMGOs, a specific inhibitor of Hippo pathway,^[16]^ XMU-MP-1, was used to treat mMGOs during the early stage of formation. XMU-MP-1 significantly promoted YAP nuclear translocation and formation of mMGOs, with larger organoids were detected in the XMU-MP-1 group on the day 7 (Figure 5A,B). In addition, XMU-MP-1 treatment increased the EdU and Ki67 positive cells (Figure 5 C,D,E,F) and the relative mRNA expression of *Ki67, Pcna, Cdk1, Cdc45* and *Ccne1* (Figure 5G), indicating the promotion of cell proliferation.

**Figure 5.**
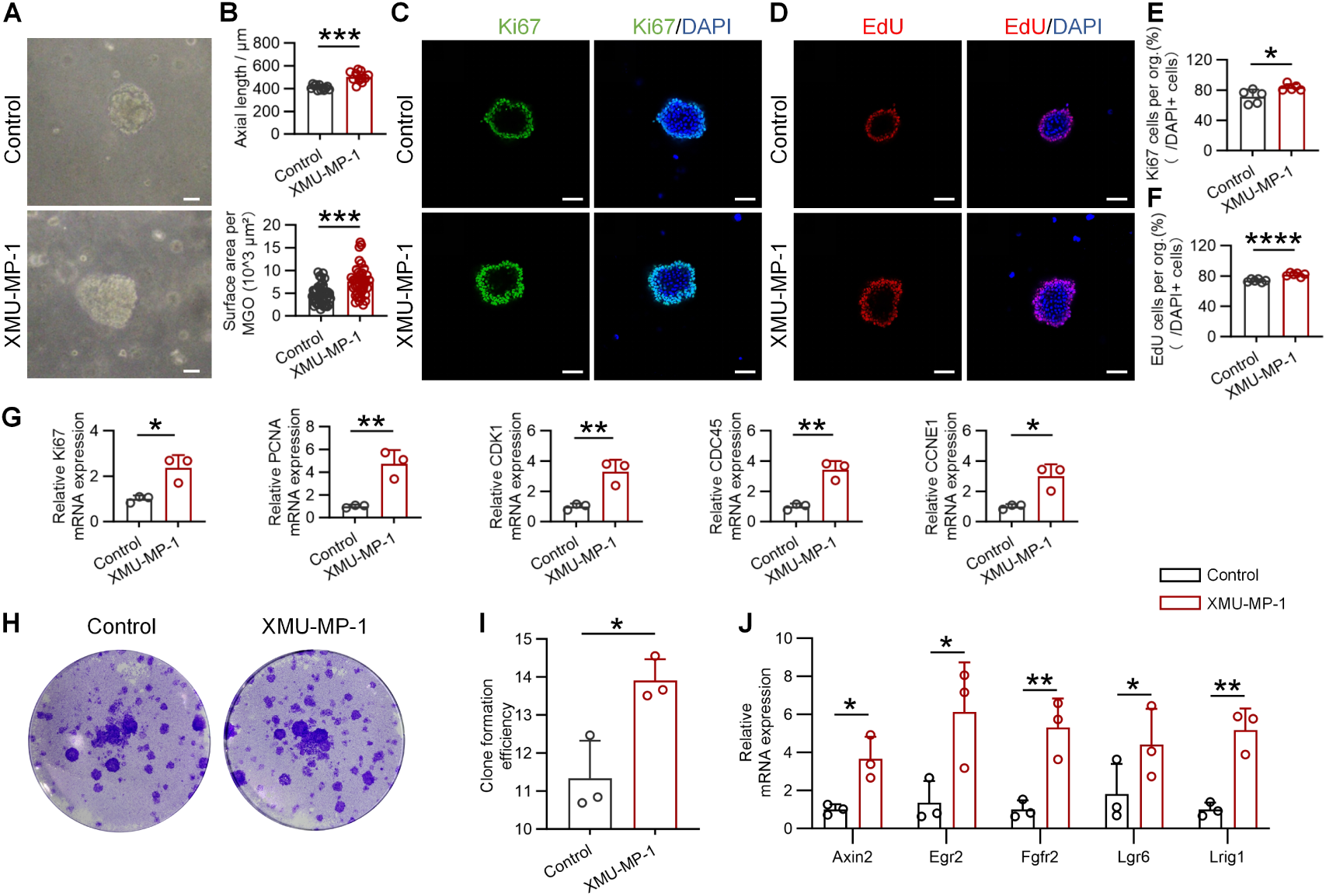
Hippo pathway inhibition enhances mMGOs formation and stem/progenitor cell clonogenicity. (A and B) Bright-field images of mMGOs treated with XMU-MP-1, with DMSO as control. The axial length (n=12) and the surface area (n=40) of mMGOs were measured using ImageJ. (C and D) Immunofluorescence of Ki67 and EdU staining in mMGOs, nuclei were counterstained with DAPI. (E and F) Percentage of Ki67 (n=5) and EdU (n=6) positive cells in (C and D). (G) Relative mRNA expression of *Ki67*, *Pcna, Cdk1*, *Cdc45* and *Ccne1* in mMGOs treated with XMU-MP-1, with DMSO as control (n=3). (H and I) Crystal violet staining of cell clones derived from mMGOs, the clone formation efficiencies were quantified by ImageJ (n=3). (J) Relative mRNA expression of *Lrig1*, *Axin2*, *Lgr6*, *Egr2* and *Fgfr2* in mMGOs treated with XMU-MP-1, with DMSO as control (n=3). Scale bars represent 100 µm in (A), 50 µm in (C) and (D).

Organoid formation requires the long-term proliferation and pluripotent differentiation potential of stem cells. Therefore, we further explored the role of Hippo pathway inhibition on the stemness of MG stem cells. Organoids were cultured under XMU-MP-1 conditions until day 7, then digested into single cells for clonal analysis. XMU-MP-1 group shows higher clone formation efficiency (Figure 5H,I). XMU-MP-1 also significantly promoted the expression of genes representing the stemness of MG stem cells, including *Lrig1, Axin2, Lgr6, Egr2* and *Fgfr2* (Figure 5J), which have been reported to be associated with MG homeostasis.^[17–19]^ These results suggesting Hippo pathway inhibition significantly promotes stemness in mMGOs.

### 2.6 YAP regulates MG homeostasis

The ultimate consequence of MGD is an imbalance in MG homeostasis, leading to glandular atrophy.^[20, 21]^ We first investigated the nuclear localization of YAP in the age-related MGD (ARMGD) mouse model (24-month-old). Compared to 8-week-old mice, YAP exhibited significant nuclear exclusion in MG cells of ARMGD mice (Figure 6A,B). Next, an *in vitro* model of MGD based on organoids were established. We treated the mMGOs with lipopolysaccharide (LPS), which has been reported to induce MGD-related proinflammatory gene expression in cultured MGC.^[22, 23]^ This *in vitro* model exhibited classic MGD-like features, including glandular atrophy (Figure 6C,D), reduced cell proliferation (Figure 6E,H), increased apoptosis (Figure 6F,I), hyperkeratinization (Figure 6G,J), and increased expression of inflammatory cytokines (Figure S3A,B,C, Supporting Information), indicating the successful establishment of the models. Notably, we also observed nuclear exclusion of YAP in this *in vitro* model (Figure 6K,L).

**Figure 6.**
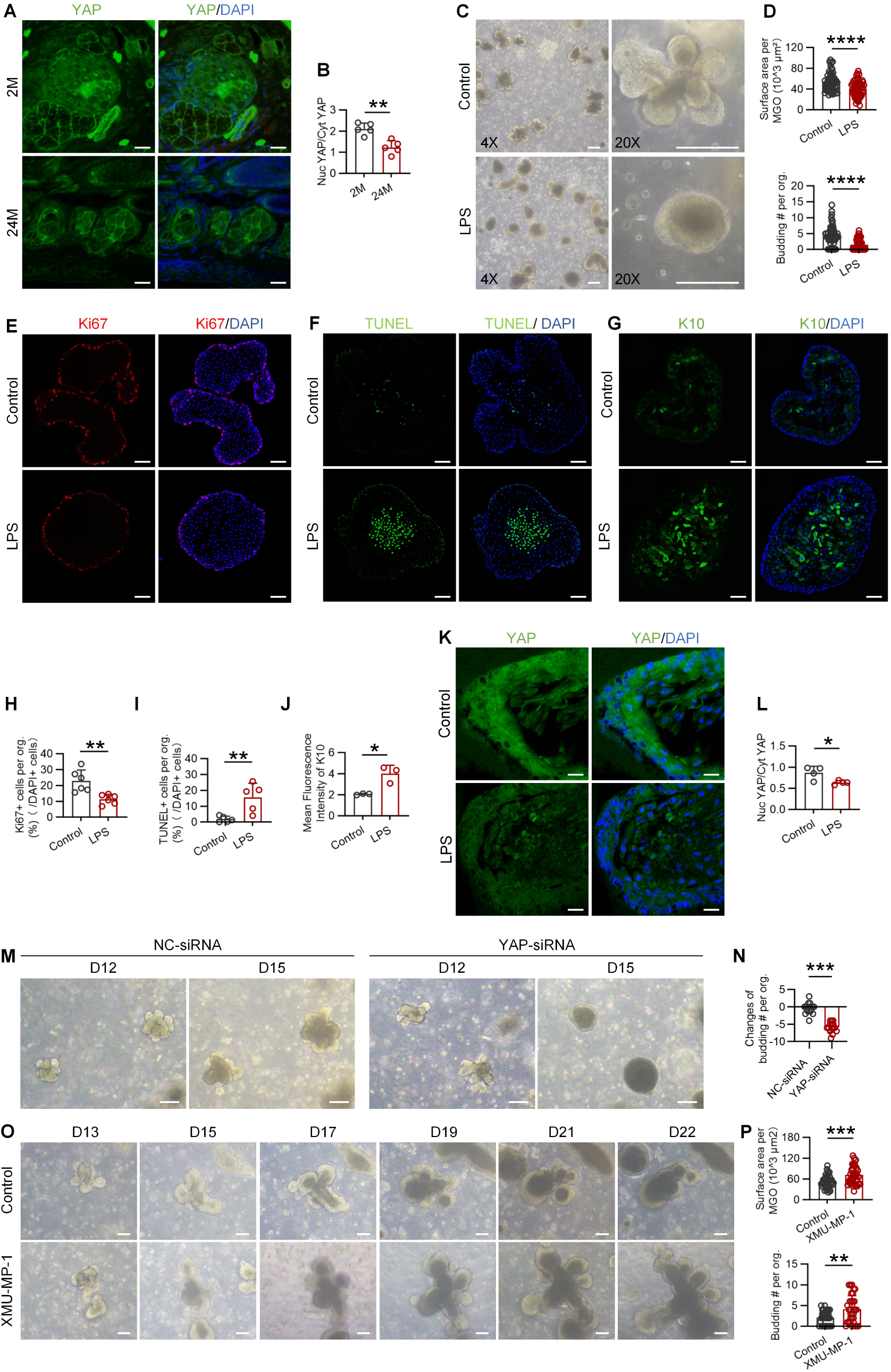
YAP sustains meibomian gland homeostasis. (A and B) Immunofluorescence of YAP in MG of 8-week-old and 24-month-old mice, nuclei were counterstained with DAPI. And the ratio of nuclear YAP to cytoplasm YAP, three cells were randomly selected for measurement and averaged in each sample (n=5). (C and D) Bright-field images of mMGOs treated with LPS, with PBS as control. Number of budding (n=62) and the surface area (n=57) of mMGOs were quantified. (E) Immunofluorescence of Ki67 in mMGOs, nuclei were counterstained with DAPI. (F) TUNEL staining of mMGOs, nuclei were counterstained with DAPI. (G) Immunofluorescence of K10 in mMGOs, nuclei were counterstained with DAPI. (H) Percentage of Ki67 positive cells in (C) (n=6). (I) The percentage of TUNEL positive cells in (D) (n=5). (J) The intensities of K10 in (E) (n=3). (K and L) Immunofluorescence of YAP in LPS-induced MGD model *in vitro*, nuclei were counterstained with DAPI. And the ratio of nuclear YAP to cytoplasm YAP (n=4). (M and N) Bright-field images of mMGOs transfected with YAP-siRNA 3 after maturation, with NC-siRNA as control. Changes of budding number were quantified (n=3). (O and P) Bright-field images of mMGOs cultured with XMU-MP-1 after maturation, with DMSO as control. Number of budding (n=33) and the surface area (n=39) of mMGOs were quantified. Scale bars represent 250 µm in (C), 100 µm in (M) and (O), 50 µm in (E), (F) and (G), 25 µm in (A) and (K).

To investigate whether glandular atrophy in MGD is associated with YAP inactivation, we knocked down *Yap1* specifically in mature mMGOs (day 10) using siRNA. Notably, organoids exhibited marked budding atrophy following *Yap1* knockdown (Figure 6M,N). In contrast, activated YAP in mature mMGOs with XMU-MP-1 effectively sustained their growth, whereas the control group exhibited significant atrophy/death by day 22 (Figure 6O,P). Taken together, YAP play a pivotal role in regulating MG homeostasis.

### 2.7 Hippo pathway inhibition promotes mMGOs regeneration following atrophy

Next, we investigate the effects of Hippo inhibition on atrophied organoids using the *in vitro* MGD model. The brightfield images reveal that mMGOs exhibit significant bud atrophy following LPS treatment, transforming from a form branched to a spherical morphology (Figure 7A). However, with XMU-MP-1 culture, the organoids began to rebud, and the branches subsequently enlarged and elongated (Figure 7A,B). YAP was observed to exhibit nuclear translocation following XMU-MP-1 treatment (Figure 7C,D). And the cell proliferation also be showed by Ki67 immunofluorescence and EdU staining (Figure 7E,F). Furthermore, K14 and K6a immunofluorescence staining confirmed the normal cells phenotype of the mMGOs (Figure 7G), and LipidTox staining indicated that the mMGOs maintained lipid synthesis function (Figure 7H).

**Figure 7.**
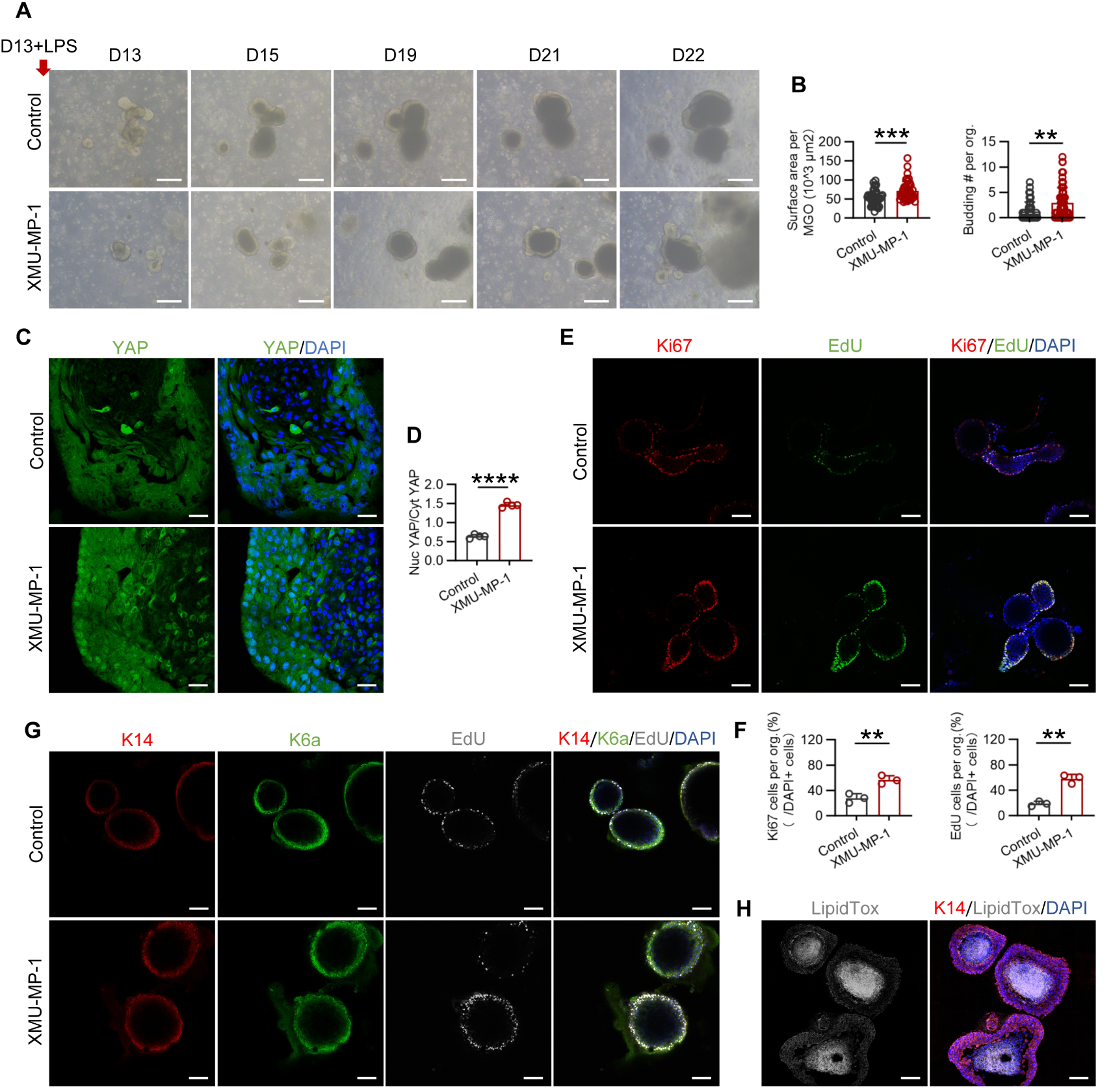
Hippo pathway inhibition promotes mMGOs regeneration following atrophy. (A and B) Bright-field images of mMGOs treated with XMU-MP-1 following LPS-induced atrophy. Number of budding (n=44) and the surface area (n=46) of mMGOs were quantified. (C and D) Immunofluorescence of YAP in mMGOs treated with XMU-MP-1 following LPS-induced atrophy, nuclei were counterstained with DAPI. And the ratio of nuclear YAP to cytoplasm YAP (n=4). (E and F) Immunofluorescence Ki67 and EdU staining in mMGOs, nuclei were counterstained with DAPI. Percentages of Ki67 and EdU positive cells were quantified (n=3). (G) Immunofluorescence of K14, K6a and EdU staining in mMGOs, nuclei were counterstained with DAPI. (H) Immunofluorescence of K14 and LipidTox saining in mMGOs. Scale bars represent 250 µm in (A), 100 µm in (H), 50 µm in (E) and (G), 25 µm in (C).

## 3. Discussion

In this study, we have developed a novel 3D organoid that faithfully mimics the postnatal development and cytodifferentiation of the murine MG. More importantly, by integrating comparative transcriptomics and functional genetics, we identified the Hippo-YAP pathway as a central regulator of MG development, epithelial homeostasis, and regeneration. Modulation of this pathway is not only required for organoid formation but is also sufficient to reactivate regenerative programs in atrophied organoids, highlighting its potential as a therapeutic target in MGD.

While recent studies have successfully established foundational MGO models for both mouse and human, enabling disease modeling with inflammatory cytokines like IL-1β and testing of known therapeutic agents (e.g., p38 MAPK inhibitors, PPARγ agonists),^[9, 10]^ these platforms primarily served to recapitulate established pathologies or validate pathways already implicated in MG biology, but their capacity to model developmental trajectories and regulatory mechanisms has remained unclear. The organoids we have established based on a novel culture strategy not only faithfully recapitulates the key biological process and temporal gene expression dynamics of postnatal MG development, but also serves as a discovery tool to identify a previously overlooked developmental regulation mechanism. We demonstrate that Hippo-YAP pathway acts as an intracellular mechanism regulating the proliferation-morphogenesis axis during organoids development. Furthermore, whereas existing MGO dysfunction models are typically used to screen for compounds that halt degeneration or alleviate symptoms, our key advancement lies in showing that pharmacological inhibition of the Hippo pathway can actively reverse and re-initiate a regenerative program in established atrophied organoids. This positions our work not only as a validation of the MGO model’s utility but as a critical step towards defining novel, mechanism-based targets for restorative therapies aimed at addressing glandular dropout in MGD.

Consistent with the well-documented pro-growth function of YAP/TAZ in other regenerating tissues,^[24–27]^ we observed nuclear translocation of YAP during both *in vivo* gland development and *in vitro* organoids formation. YAP inhibition and knockdown of *Yap1* significantly suppress MGC proliferation and mMGOs formation, indicating YAP activity as necessary condition for driving the expansion of the MGC. Conversely, activation of YAP through Hippo pathway inhibition promotes organoid growth and enhances proliferative capacity. These findings establish Hippo-YAP pathway as a critical regulatory node controlling the balance between proliferation and differentiation in MG epithelium. Future studies will be required to elucidate how YAP integrates with key differentiation regulators, such as PPAR γ, to coordinate lineage specification and lipid-producing maturation.

Notably, our study reveals a previously underappreciated role of YAP in maintaining MG homeostasis and preventing glandular atrophy. Both aging-associated MG dysfunction and LPS-induced inflammatory atrophy were associated with reduced nuclear localization of YAP, suggesting that YAP inactivation may be a common feature of MGD pathogenesis. Importantly, activation of YAP restored proliferative activity, reinitiated branching morphogenesis, and rescued lipid synthesis in MGD-like organoids. This regenerative effect is likely mediated, at least in part, through the enhancement of stem/progenitor cell function, as evidenced by increased clonogenicity and upregulation of stemness-associated genes. These results suggest that MG epithelial cells retain a degree of plasticity that can be therapeutically harnessed, and position YAP as a potential target for regenerative interventions in MGD.

To facilitate efficient formation and maturation of mMGOs, 3T3-J2 feeder cells were incorporated to provide essential niche support. This approach markedly improves organoid-forming efficiency and promotes early epithelial expansion, thereby enabling robust and reproducible organoid establishment. As a result, this system is particularly suitable for studying early developmental dynamics and morphogenetic processes of MG epithelium.

We acknowledge several limitations. First, our current model, while recapitulating epithelial components, lacks the native microenvironment, including immune cells, nerves, and the surrounding mesenchymal niche, which are known to influence MG function. Second, the functional lipidomic profile and secretory capacity of the organoids require further quantification to confirm full metabolic maturity. Third, the use of pharmacological modulators may introduce off-target effects, and future studies employing genetic lineage tracing and conditional knockout models will be necessary to validate these findings *in vivo*.

## 4. Conclusion

In summary, we present a physiologically relevant MG organoid model and use it to reveal a previously unrecognized role of Hippo-YAP pathway in regulating gland development, homeostasis, and regeneration. This work not only provides a robust experimental platform for studying MG biology and development, but also identifies a promising molecular target for regenerative therapy, paving the way for future advances in the treatment of MGD.

## 5. Methods

### Animal

All animals, regardless of gender, were housed in Xiamen University Laboratory Animal Center and maintained under a 12-h light/dark cycle, free access to water and food. Animals used in this study were treated in accordance with the ARVO Statement for the Use of Animals in Ophthalmic and Vision Research.

### Cell culture

3T3-J2 cells (OTWO, HTX2404) were cultured in Dulbecco’s Modified Eagle Medium (DMEM, Gibco, 11965092) with 10% fetal bovine serum (FBS, ABW, AB-FBS-0050) at 37℃ in 5% CO2. Before mMGOs formation and clone culture, 3T3-J2 cells were treated with mitomycin C (Sigma, M0503), which was supplemented in medium at a final concentration of 5 μg/mL for 2-4 h in an incubator to restrain the proliferation of 3T3-J2 cells.

### Mouse MG isolation and digestion

Mouse MGs were sampled under an operating microscope, followed by rinsing in Hank’s balanced salt solution (HBSS; Gibco, 14170112) supplemented with 10% penicillin-streptomycin (PS, Gibco, 15140-122) and 10% FBS at least three times. Then, the MG tissues were immersed in serum-free and hormoneenriched medium (SHEM) supplemented with 10% collagenase A and incubated at 37◦C for 12–14 h overnight. Next, mouse MG tissues were centrifuged and washed with HBSS. Single cells were further digested with 0.25% trypsin-EDTA (Gibco, 25200072) at 37◦C for 5 min. The post-digestive solution was filtered through a cell strainer with a 40 μm aperture to ensure single-cell isolation. The SHEM medium was composed of Dulbecco’s Modified Eagle Medium/Nutrient Mixture F-12 (DMEM/F12, Gibco, 11320033) supplemented with 5% FBS, 0.5% DMSO, 2 ng/mL mouse EGF (Gibco, PMG8041), 1% Insulin-Transferrin-Selenium-Ethanolamine (ITS-X, Gibco, A4000046401), 0.5 g/mL hydrocortisone (Yeasen, 54576ES03) and 1% PS. *Establishment of mMGOs:* To establish mMGOs, meibomian gland cells (MGC) isolated from mouse MG were cultured according to the protocol established in our previous study with modifications of condition.^[11]^ In brief, MGC and 3T3-J2 cells were co-cultured in 50% hemispherical matrigel (R&D, BME001-05) at cell densities of 1000 cells/well for mouse MGC, 2×10^4^ cells/well for 3T3-J2 cells. The mMGOs were cultured in SHEM medium at 37℃ in 5% CO2, with culture medium changed every other day. The Rock inhibitor Y27632 (MedChemExpress, HY-10071), was added to the SHEM medium to facilitate cell differentiation. The culture systems were observed every 2 days under a microscope. The mMGOs were photographed using phase contrast microscope (Olympus, CKX53), and the axial length and surface of mMGOs were measured by ImageJ.

### LPS induced in vitro MGD model

To induce MGD-like changes in mMGOs, 100 μg/ml LPS (Sigma, L2880) was added to the medium. After 24 h of LPS treatment, switch to normal medium and continue culturing.

### Clone culture

The clonal culture method was adapted from previous study.^[11]^ 3T3 cells and cells isolated from mMGOs were co-cultured at cell densities of less than 800 cells/cm^2^ for mMGO cells, and 5×10^4^ cells/cm^2^ for 3T3 cells. The medium is SHEM medium supplemented with 10 μM Y27632. After cell seeding, the culture systems were observed every 2 days under a microscope. On the 14th day of clonal culture, 3T3 feeder cells were digested with 0.25% trypsin-EDTA for 1 min and removed. Cell plates with MG colonies were then fixed with 4% paraformaldehyde for 10 min, followed by crystal violet staining for 30 min. Cells stained with crystal violet were photographed and analyzed later. The area and number of the MG colonies was measured using ImageJ.

### siRNA transfection

YAP expression was knocked down in mMGOs transfection with the YAP siRNA (Sangon Biotech) using RNATransMate (Sangon Biotech, E607402) following the manufacturer’s instruction, with negative control (NC) siRNA as control.

### RNA sequencing

Total RNA was extracted with Total RNA Extraction Reagent (Abclonal, RK30129) according to the manufacturer’s protocol. RNA sequencing analysis was performed on Illumina HiSeqTM 4000 system by Beijing CapitalBio Technology, China. Besides, the RNA was sheared and reverse transcribed to get cDNA used for library construction. After that, sequencing was performed on the prepared library. All the generated raw sequencing reads were filtered to get clean reads stored as FASTQ format. RSEM was used to quantify the Genes expression level (FPKM). DESeq2 package was used for comparative analysis of differences. The screening criteria for differentially expressed genes were |log2FC| >= 1 and p-Value <= 0.05. Gene Ontology (GO) and pathway annotation and enrichment analyses were based on the Gene Ontology Database (http://www.geneontology.org/) and KEGG pathway database (http://www.genome.jp/kegg/), respectively. Bioinformatic analysis was performed using Omicsmart, a dynamic real-time interactive online platform for data analysis (http://www.omicsmart.com).

### EdU labeling

EdU (APExBIO, B8337) was supplemented in the mMGOs culture medium for a final concentration of 20 mg/ml for 8 hours, then the mMGOs were collected. EdU signal was detected by a laser scanning confocal microscopy (ZEISS, LSM880) after imaging with the Click-iT EdU Alexa Fluor 647 imaging kit (Invitrogen, C10640) according to the manufacturer’ s instruction.

### Histological Analysis

Mouse eyelid tissues and mMGOs were embedded in optimal cutting temperature (OCT) compound, cut into sections with 6 µm thickness. Frozen sections were prepared for hematoxylin and eosin (H&E) staining, immunofluorescence staining, TUNEL assay, Oil Red O staining and LipidTox staining.

### Immunofluorescence staining

The mMGOs whole mount and frozen sections of mouse eyelid and mMGOs were fixed with 4% paraformaldehyde for 20 min and incubated with 2% bovine serum albumin for 60 min to reduce background noise. The mMGOs and sections were then labeled with primary rabbit antibodies for YAP1 (Abclonal, A19134), Ki67 (Abcam, ab16667), PPARγ (Abcam, ab45036), K10 (Abcam, ab76318), K14 (Abcam, ab7800), K6a (BioLegend, 905701), IL-6 (Abcam, ab7737), IL-10 (Abcam, ab133575), IL-1β (Abcam, ab9722), MMP3 (Abcam, ab52915), MMP9 (Abcam, ab38898) overnight at 4℃. After 3 washes with PBS, the samples were incubated with goat anti-rabbit lgG (Abclonal, AS053) for 120 min. The nuclei were stained with DAPI (Beyotime, C1005) for 10 min. The samples were observed under a laser scanning confocal microscopy (ZEISS, LSM880).

### TUNEL assay

TUNEL assay was performed using the DeadEnd Fluorometric TUNEL System (Promega, G3250). Frozen mMGOs sections were rehydrated and incubated with Proteinase K Tris/HCL, pH = 7.4 (10 mM) for 30 min at 37°C. After washing 3 times each with PBS for 5 min, 50 µL of TUNEL reaction mixture was added on the sections and the sections placed in the dark for 1 hour at 37°C, followed by rinsing 3 times with PBS for 5 min each. Finally, the specimens were counterstained with DAPI, mounted, and photographed with a confocal microscope (ZEISS, LSM880).

### Oil Red O staining

Frozen mouse eyelid sections were fixed in 4% paraformaldehyde for 10 min, washed in PBS for 5 min, and stained for 10 min in freshly prepared Oil Red O solution. After rinsing with PBS for 5 min, the sections were counterstained with hematoxylin and mounted in 90% glycerol.

### LipidTox staining

Frozen mMGOs sections were fixed in 4% paraformaldehyde for 10 minutes. After rinsed in PBS for 5 min, sections were incubated with the LipidTox neutral lipid stain (1:200, Thermofisher, H34475) at room temperature for 2 h, followed by counterstaining with DAPI. Sections were then imaged with a confocal microscope (ZEISS, LSM880).

### Reverse Transcription Quantitative PCR (RT-qPCR)

The total RNA was extracted using Total RNA Extraction Reagent (Abclonal, RK30129), and the cDNA was made by using the RT Master Mix for qPCR (Abclonal, RK20433). RT-qPCR was performed with the SYBR Green qPCR Mix with UDG (Abclonal, RK21219), and the primer sequences are listed in Table S1. The mixture was held at 37℃ for 2 min and then heated to 95℃ for 3 min, cycled 40 times (95℃ for 5 s, 60℃ for 30 s). Melting curves were generated by increasing the temperature from 55℃ to 95℃ in 0.5℃ increments at 10 s intervals and then visually to ensure that a single peak was present for each primer. Threshold amplification values (Ct) were assigned by the LightCycler 96 analysis software (Roche).

### Western Blot Analysis

The mMGOs were collected in RIPA lysis buffer (Solarbio, R0010) with 1% protease inhibitor (GlpBio, GK10014) and centrifuged at 12000 rcf for 15 min at 4 ◦C for protein extraction. Protein concentration was determined using Pierce BCA protein assay kit (Thermo Scientific, 23227). Protein samples were separated on 10% SDS-polyacrylamide gel by electrophoresis, then transferred to polyvinylidene difluoride membranes. The membrane was blocked with 5% milk for 2 h, then incubated in primary antibody against YAP1 (1:5000, Abclonal, A19134) and β-actin (Abcam, 8226). HRP-conjugated anti-rabbit secondary antibody (ABclonal, AS014) was then applied for 2 h after washing. The membrane was developed with ECL Chemiluminescence reagent (NCM Biotech, P10300) using ChemiDoc MP (Bio-Rad). The band intensity was semiquantified by densitometry using ImageJ software and normalized by β-actin levels.

### Statistical Analysis

Statistical analysis was conducted using the GraphPad Prism software (Graph pad 9 software Inc.). Data was presented as mean ± standard deviation. All the results were analyzed by the Kolmogrov-Smirnov normality test. Statistical comparison was conducted using the Student t-test for two groups or one-way ANOVA with multiple comparison for more than three groups of variables. p<0.05 was considered to be statistically significant.

## Ethics Statement

The animal experiments were approved by the Laboratory Animal Management and Ethics Committee of Xiamen University (XMULAC20200145).

## Acknowledgement

We thank Jingru Huang and Xiang You from Central Lab, School of Medicine, Xiamen University for technical support in confocal imaging.

## Author contributions

The experimental work in this manuscript was conducted by M.Z. and J.Z.. L.Z., R.Z. and L.S. contributed to data analysis. Manuscript writing was conducted by M.Z. and J.Z. Reviewing was conducted by W.L., Y.W. and J.B.. Y.W. and J.B. supervised the project. M.Z. and J.Z. contributed equally to this work, thus should be considered as co-first authors. All authors have reviewed and approved the manuscript.

## Data availability statement

The data supporting the findings of this study are available in the Supporting Data Values file. The raw data used in this study are available from the corresponding author upon reasonable request.

