## Supplementary figures for "Modeling Meibomian Gland Development and Dysfunction: A Mouse-Derived Organoid System Reveals Hippo-YAP as a Critical Regulator"

### Supplementary Materials

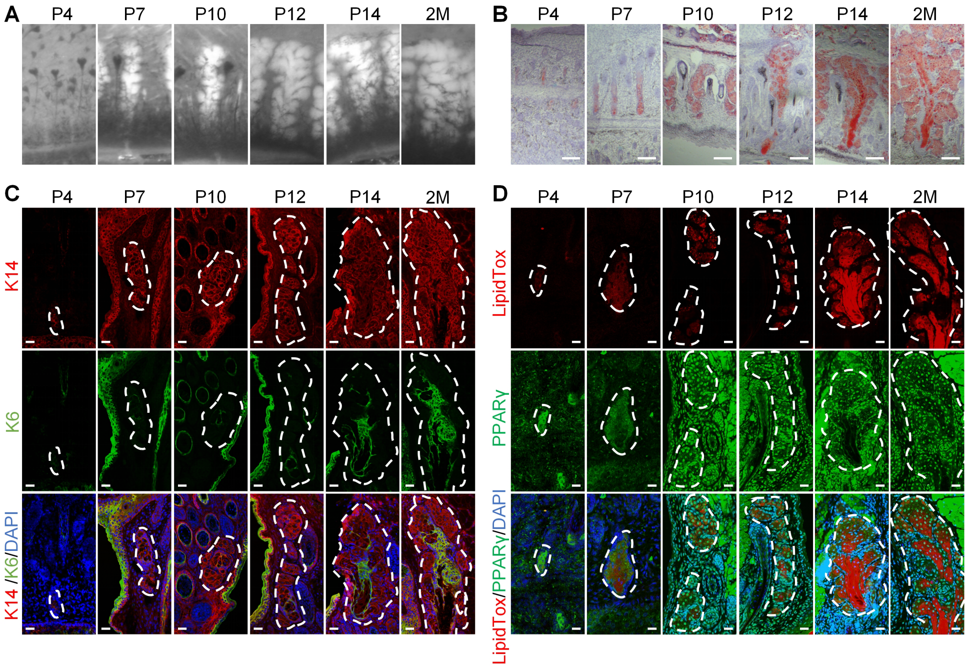


**Figure S1. Characterization of the postnatal mouse MG development.** (A) Stereomicroscope images of mouse MG during development. (B) Oil red O staining images of mouse MG during development. (C) Immunofluorescence of K14 and K6a in mouse MG, nuclei were counterstained with DAPI. (D) Immunofluorescence of PPARγ and LipidTox staining in mouse MG, nuclei were counterstained with DAPI. Scale bars represent 80 μm in (B), 25 μm in (C) and (D).

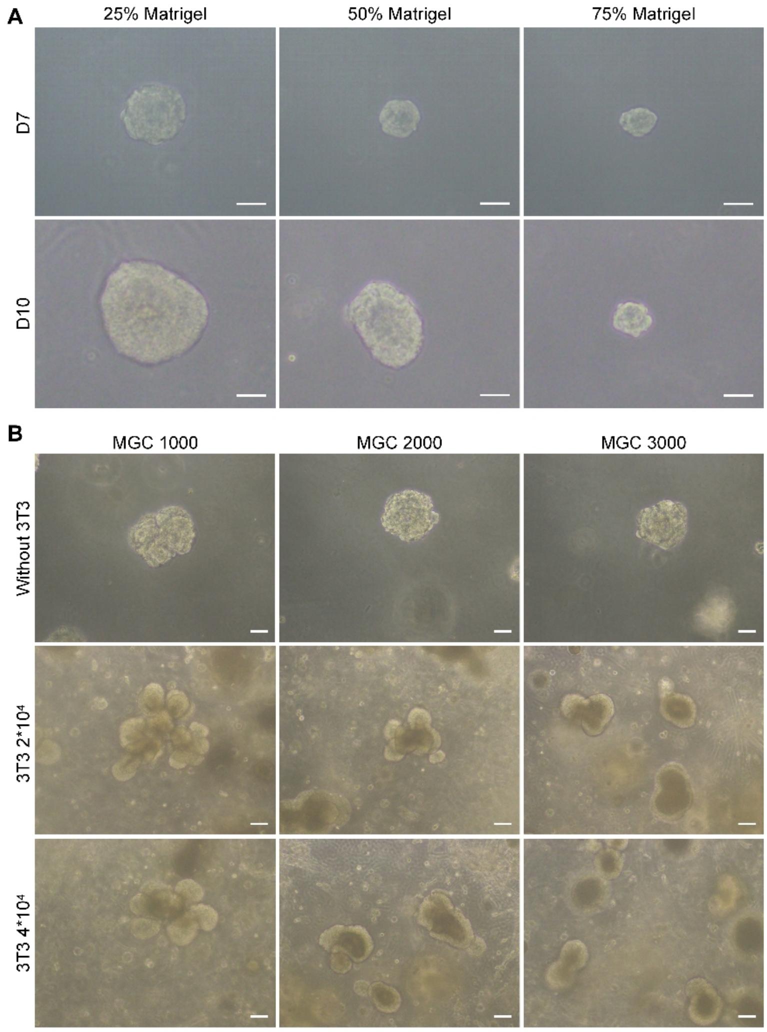


**Figure S2. Optimization of the 3D culture conditions for mMGOs formation.** (A) Bright-field images of mMGOs cultured in different concentrations of matrigel (25%, 50% and 75%). (A) Bright-field images of mMGOs formed from various combinations of meibomian gland cells and 3T3 cells. Scale bars represent 100 μm in (B), 50 μm in (A).

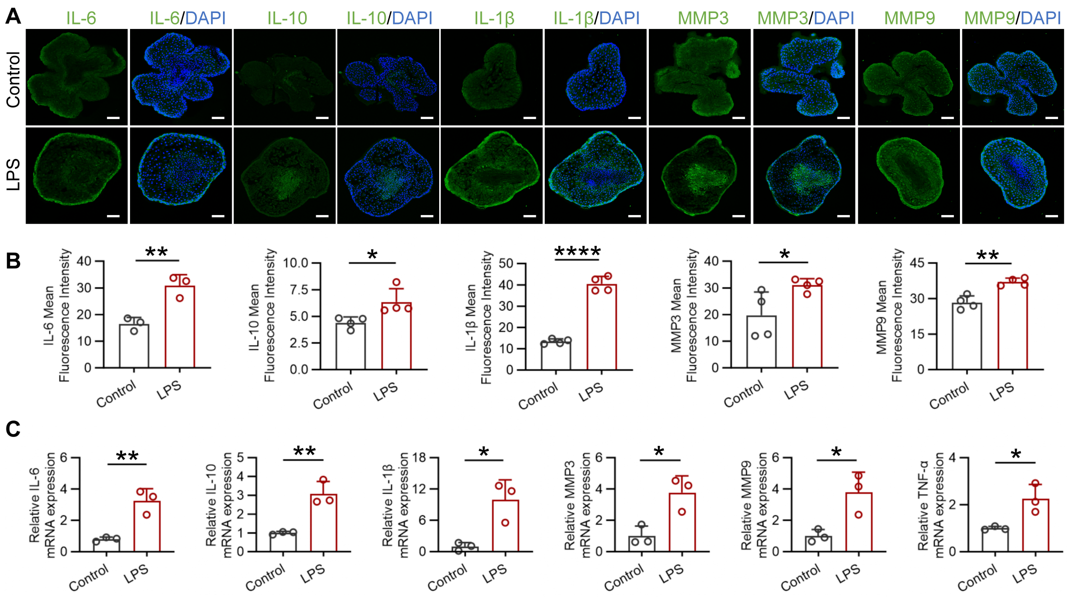


**Figure S3. LPS promotes the expression of inflammatory cytokines in mMGOs.** (A and B) Immunofluorescence of IL-6, IL-10, IL-1β, MMP3, and MMP9 in mMGOs treated with LPS, nuclei were counterstained with DAPI. The mean fluorescence intensities were quantified (n=33). (C) Relative mRNA expression of *Il-6*, *Il-10*, *Il-1β*, *Mmp3*, and *Mmp9* in mMGOs treated with LPS (n=3). Scale bars represent 50 μm in (A).

**Table S1**

| | | |
|---|---|---|
| Lrig1 | Forward | 5'-ACCACCGTAGGCATCTTCACC-3' |
| | Reverse | 5'-ACGCTGTACTCCTCGCTCTTC-3' |
| FGFR2 | Forward | 5'-GCGACTTCCAGTCAAGTGGA-3' |
| | Reverse | 5'-GTGTCCCTCTTTGAGCAGCT-3' |
| EGR2 | Forward | 5'-TGGACCACCTCTACTCTCCG-3' |
| | Reverse | 5'-GGGATGGGTAGGAAGGAGGT -3' |
| Axin2 | Forward | 5'-GCAGGCTGGCAGAGGTGTC-3' |
| | Reverse | 5'-TTGGGTTGGCGAAGGGTGAG-3' |
| CDC45 | Forward | 5'-AAGGATGGCTCAGGGACAGAC-3' |
| | Reverse | 5'-GTGGCTTGGAGGTGCTTCTTG-3' |

| | | |
|---|---|---|
| CCNE1 | Forward | 5’-GACTTACCTGAGAGATGAGCACTTTC-3’ |
| | Reverse | 5’-CGCACACCTCCATTAGCCAATC-3’ |
| CDK1 | Forward | 5’-ACGGTGTGGTGTATAAGGGTAGAC-3’ |
| | Reverse | 5’-GCACTCCTTCTTCCTCGCTTTC-3’ |
| KI67 | Forward | 5’-GCCTGCCCGACCCTACAAAATG-3’ |
| | Reverse | 5’-CTCATCTGCTGCTGCTTCTCCTTC-3’ |
| MMP3 | Forward | 5’-GTCGGGTTGGAGATGACAGGGAAG -3’ |
| | Reverse | 5’-TGAAGCCACCAACATCAGGAACAC -3’ |
| MMP9 | Forward | 5’-CGCCACCACAGCCAACTATGAC-3’ |
| | Reverse | 5’-CTGCTTGCCCAGGAAGACGAAG-3’ |
| TNF-α | Forward | 5’-GTGGTTTGTGAGTGTGAGGGTCTG-3’ |
| | Reverse | 5’-TCCCTGGGTGAGAAGCTGAAGAC-3’ |
| IL-10 | Forward | 5’-TCCCTGGGTGAGAAGCTGAAGAC-3’ |
| | Reverse | 5’-CACCTGCTCCACTGCCTTGC-3’ |
| IL-1β | Forward | 5’-GTGGTTTGTGAGTGTGAGGGTCTG-3’ |
| | Reverse | 5’-AGGTCCACGGGAAAGACACAGG-3’ |
| IL-6 | Forward | 5’-CTTCTTGGGACTGATGCTGGTGAC-3’ |
| | Reverse | 5’-AGGTCTGTTGGGAGTGGTATCCTC-3’ |
